# Systematic benchmarking of commercial workflows for isoform-resolved single-nucleus transcriptomics

**DOI:** 10.64898/2026.09.01.748518

**Authors:** Florian Köhler, Anna Delgado-Tejedor, Maik Zehnsdorf, Marisol Salgado Albarran, Laura Zalba Jadraque, Irene Gonzalez Navarrete, Luca Cozzuto, Julia Ponomarenko, Berta Fusté Pérez

## Abstract

Short-read sequencing-based single-cell transcriptomics represents the current gold standard for studying cellular transcriptomes but remains limited in its ability to resolve full-length transcript isoforms and splicing patterns. Long-read single-cell and single-nucleus RNA sequencing (LR sc/snRNA-seq) enables the transcriptome-wide characterization of full-length isoforms at cellular resolution, yet the relative performance of commercially available workflows remains insufficiently explored. Here, using nuclei extracted from a standardized multi-species benchmark sample and Oxford Nanopore Technologies long-read sequencing, we systematically benchmarked four LR snRNA-seq strategies: 10x Genomics 3’, 10x Genomics 5’, ArgenTag, and Parse Biosciences. Comparing transcriptome features qualitatively and quantitatively, as well as the concordance with matched short-read data and the ability to resolve cellular heterogeneity, we identified substantial method-specific differences in read length and yield, transcript coverage, isoform detection, and recovery of sample-specific biological information, with the 10x Genomics 3’ and 5’ assays emerging as the most balanced approaches for comprehensive isoform-resolved single-nucleus transcriptomics. Altogether, our study provides a systematic assessment of four commercially available workflows for performing LR snRNA-seq and highlights key methodological trade-offs related to distinct library preparation strategies, thus providing practical guidance for future isoform-resolved transcriptome studies at the single-nucleus level.

## Introduction

Single-cell RNA and single-nucleus RNA sequencing (sc/snRNA-seq) have transformed the study of cellular heterogeneity in recent years, yet many established methods rely on short-read sequencing as a readout. This fundamentally limits the study of transcriptome diversity, as many widely used protocols capture only the 3’ or 5’ ends of transcripts and require *in silico* reconstruction to infer transcript isoform information. As a consequence, expression analyses are often constrained to the gene level, complicating the characterization of complex transcript architectures, alternative splicing, and isoform-specific regulation, all of which can define cell states and functional phenotypes. These limitations are particularly pronounced in the nuclear compartment, where nascent transcripts, retained introns, and incompletely processed RNA species significantly expand transcriptomic diversity^1,2^. Long-read snRNA sequencing (LR snRNA-seq) represents a promising solution to overcome these bottlenecks by sequencing full-length fragments, thus enabling the unambiguous identification of transcript isoforms and providing a more comprehensive view of transcriptomic complexity at the single-nucleus level^3–5^.

Despite its potential, the adoption of LR sc/snRNA-seq as a standard approach to study the transcriptome of individual cells and nuclei has been slow, in part due to high costs, low throughput, a lack of standardized methodology, and limited read accuracy relative to short- read sequencing platforms. However, recent technological improvements have begun to help overcome these barriers. Advancements in sequencing yield and accuracy of long-read sequencing platforms such as Oxford Nanopore Technologies (ONT) and Pacific Biosciences (PacBio) have improved the experimental feasibility and financial viability of such studies. At the same time, the broader sc/snRNA-seq field has seen rapid methodological expansion, with technological innovations driving down costs, while significantly increasing the number of samples and cells that can be analyzed per experiment^6^. Specifically, continuous improvements in droplet-based approaches, such as those commercialized by 10x Genomics, together with recently developed strategies relying on combinatorial barcoding, as offered by Parse Biosciences, are driving a steady democratization of access to single-cell and single-nucleus studies. Together, these developments are bringing long-read sequencing-based transcriptomics into reach for many researchers for the first time.

In spite of these trends, the field of LR sc/snRNA-seq currently lacks an established gold-standard methodology. Multiple experimental and computational strategies have been proposed^7–13^, but systematic comparisons remain limited, making it difficult for researchers to select the best approach given their biological question and available budget. Proven sc/snRNA-seq methods, including those marketed by 10x Genomics and Parse Biosciences, excel at gene detection and cell throughput, but were originally designed and optimized for short-read sequencing platforms. Newer, native long-read solutions, such as those developed by ArgenTag, relying on instrument-free, smart barcoding for isoform-level sc/snRNA-seq studies^14^, address some of the inherent shortcomings of established methods, but lack rigorous scientific validation and benchmarking.

Here, we comprehensively benchmark four LR snRNA-seq solutions - 10x Genomics 3’ and 5’, Parse Biosciences and ArgenTag in combination with ONT long-read sequencing - with respect to their ability to characterize the full-length transcriptome at single-nucleus resolution. Using nuclei extracted from a standardized multi-species benchmark sample, we demonstrate that current LR snRNA-seq approaches can dissect cellular heterogeneity similar to established short-read sequencing-based workflows while delivering a comprehensive picture of the nuclear transcriptome, including splice isoforms. In doing so, we provide the research community with valuable insights into the practical feasibility, data quality, and comparative performance of current solutions for performing LR snRNA-seq experiments.

## Methods

### Sample preparation

A benchmark sample was generated as a mixture of three cell lines, human HEK293T embryonic kidney cells, mouse NIH-3T3 embryonic fibroblasts, and canine MDCK-2 kidney epithelial cells. Cells were cultured at 37 °C in DMEM medium supplemented with 10% Fetal Bovine Serum. On the day of harvesting, cells were washed with 1X Phosphate-buffered Saline (PBS), trypsinized, pelleted, and counted using a Countess II automated cell counter (AMQAX1000, Invitrogen by Thermo Fisher Scientific, Waltham, Massachusetts, USA). The cells were then pooled at a ratio of 40% human, 40% mouse and 20% dog, respectively, aliquoted at 500,000 cells per aliquot and stored in liquid nitrogen for future use.

On the day of each experiment, aliquots were retrieved from the liquid nitrogen storage and incubated at room temperature (RT) until fully thawed. Each aliquot was processed independently following the same procedure. Aliquots were first transferred to a 15-ml Falcon tube using a 1,000-μl tip without mixing. Then, 1,000 μl of pre-warmed (37 °C) Hibernate A medium (A1247501, Gibco by Thermo Fisher Scientific, Waltham, Massachusetts, USA) was added dropwise under gentle swirling, before letting the sample rest for 1 minute. Subsequently, an additional 2,000 μl of pre-warmed Hibernate A medium was added dropwise under gentle swirling. After letting the sample rest for another minute, 3,000 μl pre-warmed Hibernate A medium was added dropwise before gently inverting the 15-ml Falcon tube six times. The sample was incubated at RT again for 1 minute, 5,000 μl of pre-warmed Hibernate A medium was added dropwise, and the Falcon tube was gently inverted six times. After letting the sample rest for 1 minute, cells were pelleted by centrifugation at 350 g for 5 minutes at 4 °C. The supernatant was carefully removed, and the pellet was resuspended in 1,500 μl 1X PBS with 0.04% (w/v) Bovine Serum Albumin (BSA) by gentle pipetting up and down several times. After pooling the resuspended pellets from both aliquots, cell numbers and their viability were assessed using a Countess II automated cell counter. The sample was then stored on ice until nuclei isolation.

For nuclei extraction, the sample was centrifuged at 300 g for 5 minutes at 4 °C followed by removal of the supernatant. The cell pellet was then resuspended in 400 μl cold Lysis Buffer (10 mM Tris-HCl pH 7.5, 10 mM NaCl, 3 mM MgCl2, 0.025% (v/v) IGEPAL, 1X PBS) using a wide- bore pipette tip and gentle pipette-mixing five times. Cells were then lysed for 10 minutes on ice. Subsequently, 1,600 μl Nuclei Wash and Resuspension Buffer (1% (w/v) BSA, 0.2 U/μl Protector RNase Inhibitor (3335402001, Sigma-Aldrich, Darmstadt, Germany), 1X PBS) was added, followed by gentle pipette-mixing five times and centrifugation at 400 g for 10 minutes at 4 °C. After supernatant removal, nuclei were washed twice by resuspending in 2 ml Nuclei Wash and Resuspension Buffer, gentle pipette-mixing five times and centrifuging at 400 g for 10 minutes at 4 °C. After the third wash, nuclei were resuspended in 500 μl Nuclei Wash and Resuspension Buffer, passed through a 40-μm Scienceware Flowmi Cell Strainer (H13680- 0040, SP Bel-Art, Wayne, New Jersey, USA) into a new 15-ml Falcon tube, and counted once more using a Countess II automated cell counter.

## Library preparation

### 10x Genomics 3’ and 10x Genomics 5’ snRNA-seq

10x Genomics samples were prepared using the Chromium GEM-X Single Cell 3’ Gene Expression Reagent Kit v4 (PN-1000686, 10x Genomics, Pleasanton, California, USA) and the Chromium GEM-X Single Cell 5’ Gene Expression Reagent Kit v3 (PN-1000695, 10x Genomics, Pleasanton, California, USA) on a Chromium X Series instrument (PN-1000326, Model 00013- 03, 10x Genomics, Pleasanton, California, USA) following the manufacturer’s instructions with a targeted recovery of 10,000 nuclei per sample.

For Illumina short-read sequencing, the instructions laid out in the manufacturer’s protocols CG000731 (Rev B, August 2024) and CG000733 (Rev A, March 2024) were followed until the library stage.

For ONT long-read sequencing, 10x Genomics 3’ and 5’ cDNA libraries were prepared with the Ligation Sequencing Kit V14 (SQK-LSK114, ONT, Oxford, UK) following the two ONT protocols *Ligation sequencing V14 — single-cell transcriptomics with 3’ cDNA prepared using 10X Genomics on PromethION (SQK-LSK114)* [SST_9198_v114_revJ_13Nov2024] and *Ligation sequencing V14 — single-cell transcriptomics with 5’ cDNA prepared using 10X Genomics on PromethION (SQK-LSK114)* [SST_9204_v114_revG_06Mar2024], respectively. For both assays, 10 ng of unfragmented, full-length cDNA transcripts were used as starting material, with 200 fmol of cDNA amplicons taken forward into the end-prep step. At the subsequent adapter ligation step, the Quick T4 DNA Ligase from the NEBNext Quick Ligation Module (E6056S, New England Biolabs, Ipswich, Massachusetts, USA) was used. During the following bead clean-up, the AMPure XP bead ratio was increased from 0.4X to 0.5X to maximize the recovery of the cDNA library.

### · Parse Biosciences snRNA-seq

Parse Biosciences samples were prepared using the Evercode WT Mini v3 Single Cell Whole Transcriptome Kit (ECWT3100, Parse Biosciences, Seattle, Washington, USA) following the manufacturer’s instructions with a targeted recovery of 5,000-10,000 nuclei per sample.

For Illumina short-read sequencing, the instructions laid out in the manufacturer’s protocol UMWT3100 (v1.5, November 2024) were followed until the library stage.

For ONT long-read sequencing, Parse Biosciences cDNA library pools were prepared with the Ligation Sequencing Kit V14 (SQK-LSK114, ONT, Oxford, UK) following the ONT protocol *Ligation sequencing V14 — Direct cDNA sequencing (SQK-LSK114)* [DCS_9187_v114_revJ_12Dec2024] for PromethION, starting at the cDNA repair and end-prep step. Specifically, 100 fmol from each of the two generated cDNA sub-libraries were pooled together, adjusted with nuclease-free water to a final volume of 20 μl, and subsequently end-repaired and dA-tailed using the NEBNext Ultra II End Repair/dA-Tailing Module (E7546S, New England Biolabs, Ipswich, Massachusetts, USA) following the ONT protocol instructions. The Quick T4 DNA Ligase from the NEBNext Quick Ligation Module (E6056S, New England Biolabs, Ipswich, Massachusetts, USA) was used for the subsequent ligation of the Ligation Adapter (LA) to the repaired and end-prepped cDNA amplicons. During the following bead clean-up, the AMPure XP bead ratio was increased from 0.4X to 0.5X to maximize the recovery of the cDNA library pools.

### · ArgenTag snRNA-seq

ArgenTag samples were prepared using the Single-Nucleus RNA Library Kit for Long-Read Sequencing v1 (AT002-SD2, ArgenTag Corporation, New York City, New York, USA) targeting 10,000 captured nuclei per sample following the manufacturer’s instructions (User manual v1.0, June 2025).

For Illumina short-read sequencing, ArgenTag libraries were prepared using the NEBNext Ultra II DNA Library Prep Kit for Illumina (E7645S, New England Biolabs, Ipswich, Massachusetts, USA) according to the manufacturer’s instructions (User manual v7.0, September 2022). Briefly, 5 ng of cDNA was first fragmented to a size range of 200–500 bp on a Covaris S2 S-Series Focused-Ultrasonicator (600028, Covaris, Woburn, Massachusetts, USA), then end-repaired and dA-tailed, and ligated to the NEBNext hairpin loop adapter, which was subsequently cleaved by a USER enzyme treatment. Library amplification was conducted for 8 PCR cycles using the NEBNext Multiplex Oligos for Illumina – 96 Unique Dual Index Primer Pairs (E6440S, New England Biolabs, Ipswich, Massachusetts, USA). All bead purification steps were performed with AMPure XP beads (A63882, Beckman Coulter, Brea, California, USA).

For ONT long-read sequencing, ArgenTag libraries were prepared with the Ligation Sequencing Kit V14 (SQK-LSK114, ONT, Oxford, UK). As with the Parse Biosciences scRNA-seq library preparations, the ONT protocol *Ligation sequencing V14 — Direct cDNA sequencing (SQK- LSK114)* [DCS_9187_v114_revJ_12Dec2024] for PromethION was applied, starting at the cDNA repair and end-prep step. Briefly, 150 fmol of generated cDNA was adjusted with nuclease-free water to a final volume of 20 μl and subsequently end-repaired and dA-tailed using the NEBNext Ultra II End Repair/dA-Tailing Module (E7546S, New England Biolabs, Ipswich, Massachusetts, USA) according to the manufacturer’s instructions, with the exception that both incubation times at 20 °C and 65 °C were each extended from 5 to 15 minutes. At the subsequent adapter ligation step, the Quick T4 DNA Ligase from the NEBNext Quick Ligation Module (E6056S, New England Biolabs, Ipswich, Massachusetts, USA) was used. During the following bead clean-up, the AMPure XP bead ratio was increased from 0.4X to 0.5X to maximize the recovery of the cDNA library.

### · Quality control of 10x Genomics 3’, 10x Genomics 5’, Parse Biosciences and ArgenTag snRNA-seq libraries

The final short-read sequencing libraries were analyzed for their size distribution on an Agilent 2100 Bioanalyzer system (G2939BA, Agilent Technologies, Santa Clara, California, USA) with the Agilent High Sensitivity DNA Kit (5067-4626, Agilent Technologies, Santa Clara, California, USA), quantified via qPCR using the KAPA Library Quantification Kit KK4873 for Illumina sequencing (07960336001, Roche Sequencing Solutions, Pleasanton, California, USA), and stored at -20 °C until sequencing.

The final long-read sequencing libraries were quantified on a Qubit 2.0 fluorometer (Q32866, Invitrogen by Thermo Fisher Scientific, Waltham, Massachusetts, USA) with the Qubit dsDNA HS Assay Kit (Q32854, Invitrogen by Thermo Fisher Scientific, Waltham, Massachusetts, USA) and stored on ice until sequencing.

## Sequencing

### · Short-read sequencing

Aiming for a minimum of 20,000 reads per cell, all short-read libraries were sequenced on an Illumina platform as follows: 10x Genomics Chromium GEM-X 3’ snRNA short-read sequencing was performed on a NextSeq 2000 Sequencing System (20038897, Illumina, San Diego, California, USA) using the NextSeq 1000/2000 P2 Reagent Kit v3 – 100 Cycles (20046811, Illumina, San Diego, California, USA) utilizing standard sequencing-by-synthesis (SBS) chemistry with the following configuration (R1-I1-I2-R2): 28-10-10-90, in accordance with the 10x Genomics User Guide (CG000731, Rev B, August 2024).

10x Genomics Chromium GEM-X 5’ snRNA short-read sequencing was performed on a NovaSeq 6000 Sequencing System (20012850, Illumina, San Diego, California, USA) utilizing standard SBS chemistry with the following configuration (R1-I1-I2-R2): 28-10-10-90, in accordance with the 10x Genomics User Guide (CG000733, Rev A, March 2024).

Parse Biosciences Evercode WT Mini v3 snRNA short-read sequencing was performed on a NextSeq 2000 Sequencing System (20038897, Illumina, San Diego, California, USA) using the NextSeq 1000/2000 P2 Reagent Kit v3 – 100 Cycles (20046811, Illumina, San Diego, California, USA) utilizing standard SBS chemistry with the following configuration (R1-I1-I2-R2): 64-8-8- 58, in accordance with the Parse Biosciences User Guide (UMWT3100, v1.5, November 2024). ArgenTag snRNA short-read sequencing was performed on a NovaSeq X Plus Sequencing System (20084804, Illumina, San Diego, California, USA) utilizing XLEAP-SBS chemistry with a symmetrical read length of 151 cycles for Read 1 (R1), 10 cycles each for the i7 and i5 index reads, and 151 cycles for Read 2 (R2).

### · Long-read sequencing

Long-read sequencing was performed on a PromethION 2 (P2) Solo sequencer (PRO-SEQ002, ONT, Oxford, UK) connected to an HP Z8 Fury G5 workstation (66A36AV, HP, Palo Alto, California, USA) using PromethION DNA flow cells with R10-series (R10.4.1) nanopores (FLO- PRO114M, ONT, Oxford, UK). Sequencing runs were configured via MinKNOW (v25.05.14), with run parameters set to: (i) run chemistry: flow cell type=FLO-PRO114M, kit type=SQK- LSK114; (ii) run settings: run limit=72 hours, pore scan frequency=1.5 hours, reserved pores=off, real-time basecalling=high-accuracy (HAC) model, modified basecalling=off, minimum Q score (pass)=10, minimum filter read length (pass)=200 bases. Depending on the expected final cDNA library average length, sequencing libraries were split into two aliquots with molar amounts of 35 fmol (ArgenTag), 40 fmol (10x Genomics 3’ and 5’ assays), or 75 fmol (Parse Biosciences) each. ONT PromethION DNA flow cells were subsequently primed and loaded following the manufacturer’s instructions outlined in the corresponding ONT protocols for: (i) 10x Genomics 3’ cDNA libraries: SST_9198_v114_revJ_13Nov2024; (ii) 10x Genomics 5’ cDNA libraries: SST_9204_v114_revG_06Mar2024; and (iii) Parse Biosciences and ArgenTag cDNA libraries: DCS_9187_v114_revJ_12Dec2024. Each library was sequenced on two ONT PromethION DNA flow cells for a total of 72 hours per flow cell without performing a flow-cell wash or subsequent reloading.

## Data processing

### · Long-read data processing

POD5 files were basecalled using Dorado (v0.8.0) with the super-accuracy (SUP) model (dna_r10.4.1_e8.2_400bps_sup@v5.0.0), aiming to achieve the highest possible accuracy. No demultiplexing nor trimming was performed in this step. Basecalled data were then input into a custom Nextflow pipeline, which is based on scnanoseq of nf-core (v1.2.1)^15^. Initially, raw FASTQ files generated by Dorado (v0.8.0) were used to identify, extract, and correct cell barcodes (CB), as well as UMIs. Demultiplexing was performed in a platform-dependent manner. For 10x Genomics 3’ and 5’ datasets, BLAZE (v2.5.1)^16^ was used with the expected number of cells set to 10,000 and the specific options --full-bc-whitelist 3M-3pgex-may- 2023_TRU.txt.zip --kit-version 3v4 and --full-bc-whitelist 3M-5pgex-jan-2023.txt.zip --kit-version 5v3, respectively. For ArgenTag data, chimeric reads were split using a custom Bash script (split.sh) provided by the manufacturer. Then, the output was demultiplexed by taggy_demux (v1.0.0) (https://github.com/argentagsw/taggy_demux) using the options –out- fmt=’flames’ –trim-TSO –preserve –trim-poly ‘normal’ –orient ‘sense’ –keep-failed. For Parse Biosciences data, reads were first split into R1 and R2 to mimic the short-read structure using a modified custom Python script provided by Parse Biosciences with the options –chemistry v3 –l1dist 3 –l2dist 2, and then input into split-pipe (v1.3.1) (https://support.parsebiosciences.com/hc/en-us/article_attachments/31747222032276) with the options –mode pre –one_step –chemistry v3 –kit WT_mini.

Subsequently, the tagged files were processed in a uniform manner. First, alignment was performed using minimap2 (v2.28)^17^ with the options –MD -ax splice -I12g -un -k14 – secondary=no, against the combined genomes of human (Homo_sapiens.GRCh38.dna.fa), mouse (Mus_musculus.GRCm39.dna.fa), and dog (Canis_lupus_familiaris.ROS_Cfam_1.0.dna.fa) from Ensembl (v111)^18^. Second, the obtained BAM files were processed to extract the corrected CB and UMI of each read ID into specific tags (CB and UB) using custom Python scripts. Lastly, aligned and tagged reads were deduplicated using umi-tools (v1.1.5)^19^ with the options –random-seed=100 –per-cell. For ArgenTag and Parse Biosciences, the additional arguments –extract-umi-method=tag –cell- tag=CB –umi-tag=UB –method directional were used. Finally, deduplicated BAM files were used as input for IsoQuant (v3.10)^20^ with the options –stranded none –gene_quantification all –transcript_quantification all – splice_correction_strategy default_ont –model_construction_strategy sensitive_ont – data_type nanopore –counts_format mtx –normalization_method none to obtain the gene and isoform quantifications at cell level. IsoQuant generates two types of output files that were processed differently. The GTF from the captured transcriptome was analyzed by SQANTI3 (v5.5.4)^21^ with the options –report both to assess its quality and characteristics, and the count matrix (.mtx file) with barcodes and features (.tsv files) were input into a custom R script that runs an initial QC analysis (number of cells, number of features), as well as generating the gene and isoform matrix in .h5ad and .h5 Seurat formats.

### · Short-read data processing

Parse Biosciences data were processed using split-pipe (v1.3.1), while 10x Genomics data were processed using CellRanger (v8.0.1). For ArgenTag short-read data, raw sequencing reads in FASTQ format were demultiplexed using taggy_demux (v3.3.2) (https://github.com/argentagsw/taggy_demux) with the options --presets=illu+v1 and -- mate-file pointing to the R2 read file. Cell barcodes were subsequently extracted from the resulting FASTQ pair files and converted into 10x Genomics-compatible barcode sequences (i.e., nucleotide sequences) by mapping against the 10x Genomics whitelist *3M-february- 2018.txt.gz* using a custom Python script provided by ArgenTag. The resulting R1 and R2 files were then processed using CellRanger (v9.0.1) with the parameter --chemistry=SC3Pv3 to perform alignment, barcode correction, and gene expression quantification.

## Data analysis

### · Read retention

Input reads were defined as raw, unfiltered sequencing reads. Read retention was calculated as the percentage of reads remaining at each processing step relative to the raw reads. These steps included cell barcode identification (tagged), read alignment by minimap2 (mapped), deduplication using UMI-tools (dedup), and assignment to genes and isoforms using IsoQuant (v3.10)^20^ (Suppl. Tab. 2). For each single-cell sequencing platform, the median read retention across replicates (n=3) was calculated and plotted using a custom python script, with bars representing medians and error bars indicating standard deviations.

### · Metagene coverage

Metagene coverage profiles were generated from deduplicated BAM files using deeptools (v3.5.4)^22^. Alignments with MAPQ scores lower than 10 were excluded, and transcripts shorter than 350 bp were filtered to achieve sufficient resolution for coverage analysis. The coverage across the gene body – from TSS to TES – was computed separately for the top 10% longest and shortest transcripts, respectively. All three replicates per sequencing platform were included in the analysis to obtain comprehensive coverage profiles. Coverage metrics were calculated and visualized for each platform independently, enabling platform-specific assessment of gene body coverage distribution. Lastly, the profiles obtained per platform were overlapped to allow for an easier visual comparison.

### · Base distribution across feature types

The proportion of bases mapping to each RNA feature category (coding, intronic, UTR, intergenic, and non-aligned) was assessed for each sample using CollectRnaSeqMetrics of the Picard toolkit (v3.4.0) (https://broadinstitute.github.io/picard/). Median values were then calculated per platform (10x Genomics 3’, 10x Genomics 5’, ArgenTag, and Parse Biosciences) and visualized with error bars representing standard deviations.

### · Isoform analysis at bulk level (SQANTI3)

SQANTI3 results (v5.5.4)^21^ were analyzed to characterize the transcriptome captured by each single cell platform. First, the median number of genes and isoforms captured was calculated. Second, isoforms were classified into four structural categories: (i) Full Splice Match (FSM) transcripts, which match a reference transcript at all splice junctions; (ii) Incomplete Splice Match (ISM) transcripts, which match consecutive but not all splice junctions of the reference transcripts, and two novel transcript categories; (iii) Novel in Catalog (NIC) transcripts, which contain new combinations of already annotated splice junctions or novel splice junctions formed from already annotated donors and acceptors; and (iv) Novel Not in Catalog (NNC) transcripts, which use novel donors and/or acceptors^21^. Moreover, the relationship between genes and isoforms in novel versus annotated for each category was assessed. For all these metrics, the median value was calculated across replicates, with error bars representing standard deviations.

SQANTI3 also characterized all splice junctions captured by each platform, which can be classified as canonical (consensus GT-AG splice sites) or non-canonical (all other configurations), and additionally categorized as known (matching annotated junctions) or novel (not previously annotated). Junctions consistently detected across all three replicates per platform were retained for analysis to ensure robust junction characterization. For all these metrics, the median value was calculated across replicates, with error bars representing standard deviations.

### · Gene and isoform-cell matrixes preprocessing

The gene and isoform-cell matrixes generated by IsoQuant were further analyzed using a custom pipeline based on Python (v3.10) and the Scanpy (v1.10.1)^23^ package.

Feature identifiers were extracted from the raw matrices and cross-referenced against the Ensembl BioMart database (October 2024)^18^ to retrieve associated biological metadata, including gene and transcript names, Ensembl IDs, and chromosomal locations. Features located in the chromosome MT were labeled as mitochondrial, and features, whose gene name matched the pattern RP[SL]Δ or MRP[SL]Δ, as ribosomal.

For the non-10x Genomics platforms, demultiplexing algorithms do not perform cell calling, whereas BLAZE (v2.5.1)^16^ does. Therefore, automated cell calling was performed for ArgenTag and Parse Biosciences gene matrixes using a knee-point detection algorithm applied to the barcode rank plot, followed by threshold-based filtering at 5% of the inflection point. The true cell barcodes were extracted from the gene matrixes, and those were used to subset the isoform ones (Suppl. Tab. 3). Then, all matrixes were processed in the same manner. Nuclei were assigned to the dominant species, if a species accounted for ≥80% of the nucleus’ total counts, and nuclei with mixed-species dominance (no single species exceeding the threshold) were flagged as mixed. Quality control metrics were calculated including total counts per cell, number of genes per cell, and percentage of mitochondrial/ribosomal counts. Finally, matrixes belonging to the same single-cell platform (10x Genomics 3’, 10x Genomics 5’, ArgenTag, and Parse Biosciences) were merged using outer join on feature annotations, with duplicate entries retained only once.

### · QC-based filtering for gene and isoform-cell matrixes

Quality control-based filtering was applied to annotated count matrices to remove low-quality nuclei and genes. Features (genes/transcripts) were initially filtered, retaining only those expressed in a minimum of 15 nuclei. Nuclei were retained if they simultaneously met the following criteria: (i) ≥1,000 genes detected (minimum complexity threshold); (ii) ≤5,000 genes detected (upper complexity threshold to exclude potential doublets); (iii) ≤15% mitochondrial counts (to exclude dying or damaged cells); and (iv) ≤20,000 total counts (to exclude doublets with inflated expression). For gene-level data, doublet detection was performed using sc.pp.scrublet (Scanpy, v1.10.1)^24^ and doublet-predicted nuclei were removed from downstream analysis (Suppl. Tab. 4). Quality control metrics were visualized before and after filtering, with plots displaying the distribution of genes per nucleus, total counts per nucleus, mitochondrial percentage, and ribosomal percentage for each sample. Finally, nuclei were assigned to the dominant species if that species accounted for ≥80% of the nucleus’ total counts. Nuclei with mixed-species dominance (no single species exceeding the threshold) were flagged as mixed (Suppl. Tab. 5). Filtered matrices were saved for subsequent analysis.

### · Dimensionality reduction, batch correction and clustering

Filtered count matrices underwent log-normalization, with counts per nucleus normalized to 10,000 and log-transformed. Raw counts were preserved in a separate layer for downstream differential expression analysis. Highly variable genes (HVGs) were identified while excluding mitochondrial genes, selecting the top 10% of genes ranked by variance using the Seurat v3 method with sample as a batch key. Principal component analysis (PCA) was performed on the HVG set retaining the top 100 components. Batch effects across samples were corrected using sc.external.pp.harmony_integrate (Scanpy, v1.10.1)^23^, which iteratively adjusted the PCA representation to remove sample-specific variations while preserving biological differences. Corrected PCA coordinates were used to construct k-nearest neighbor (KNN) graphs (k=15) and to generate UMAP plots for visualization. Leiden clustering was applied at six resolutions (0.05, 0.1, 0.3, 0.5, 1.0, 1.5) to identify clusters at varying granularities. Silhouette scoring was computed for each resolution to assess the cluster quality by selecting the optimal resolution, defined as the resolution with the highest average silhouette coefficient.

### · Cell-cycle scoring

The cell cycle phase was assessed for all nuclei at isoform level using a set of mouse marker genes, which are orthologous to the human genes described by Tirosh et al.^25^ The reference gene set consisted of 43 S-phase genes and 54 G2/M-phase genes identified by Ensembl IDs. For each sample, all expressed transcripts belonging to the set of genes were used for scoring. Briefly, expression data were first normalized to 10,000 counts per nucleus and log-transformed. Next, cell cycle scores were computed using Scanpy (v1.10.1)^23^, calculating a per- cell signature score based on the mean expression of S-phase and G2/M-phase genes. Two scores, S_score and G2M_score, were assigned to each nucleus. Phase assignment thresholds were calculated as follows: (i) if the median score was below zero, the threshold was set to zero plus the standard deviation; otherwise, (ii) the threshold was set to the median plus the standard deviation. Nuclei were assigned to the S phase if the S_score exceeded its calculated threshold and was greater than the G2M_score. Conversely, nuclei were assigned to the G2/M phase if the G2M_score exceeded its calculated threshold and was greater than the S_score. All remaining nuclei were classified into the G1 phase. Results were visualized as UMAP plots, colored by the respective S_score and G2/M score, as well as by the assigned phase.

### · Downsampling at nuclei and sequencing depth

Downsampling analysis was performed starting from the pooled isoform-by-cell matrices generated for each platform. In brief, 3,000 nuclei were randomly sampled from each initial pooled matrix, excluding nuclei previously classified as mixed and without constraining the relative proportions of human, mouse, and dog nuclei. Because this sampling is stochastic, the procedure was repeated three times to account for the variability introduced by the random selection, yielding three nucleus-level downsampled matrices per platform.

These matrices were then further downsampled to a range of sequencing depths per nucleus (3,000; 2,000; 1,000; 500; and 100 UMIs/nucleus). Given the stochastic nature of this step, it was likewise repeated three times for each input matrix. In total, nine matrices were obtained per platform and per sequencing depth.

### · Per-cell sequencing saturation

For each platform, 3,000 nuclei with at least 3,000 UMIs/nucleus were randomly sampled. Sampling depths were then set as a series of evenly spaced values between zero and the minimal UMIs across platforms. For each nucleus, UMIs were randomly subsampled to each sampling depth, and the number of unique features detected was counted at each depth. Saturation was then calculated as 1 - (number of features detected / UMIs sampled). Per- platform curves show the mean saturation across the sampled nuclei, with bands denoting ±1 standard deviation. The reported saturation value corresponds to the deepest sampling depth. This analysis was done both at isoform and at gene level.

Note that this rarefaction-based metric is computed on deduplicated UMIs from the count matrix and quantifies how many additional features each further UMI reveals. It therefore differs from PCR-duplicate–based sequencing saturation (1 − unique reads / total reads), which is calculated from raw alignments prior to deduplication.

## · Feature replicability

For each platform, features were first compared within each set of 3,000 nuclei. The three subsampled matrices generated from the same 3,000 nuclei at 3,000 UMIs/nuclei were intersected, retaining the features detected in all three. The three resulting sets per platform were then combined by union. Finally, these per-platform sets were compared across platforms to identify features consistently detected in every platform, and all overlaps were visualized using UpSet plots (v0.9.0; doi.org/10.1109/TVCG.2014.2346248). This analysis was performed at both gene and transcript level.

### · Short- and long-read gene count matrixes comparison

Matched short-read and long-read datasets from the same biological samples were analyzed. First, cell barcodes were extracted from both modalities, and datasets were filtered to retain only nuclei with matching barcodes. For the ArgenTag platform, cell barcodes in the short-read dataset are initially encoded as nucleotide sequences and require translation to numeric IDs to enable barcode matching. This operation was executed using a custom Python script that takes as input the mapping between the nucleotide sequences and the numeric IDs provided by ArgenTag. Genes with zero total counts were removed per modality. For each modality, raw count matrices were aggregated by summing counts across all filtered cells per gene, producing one expression profile per modality. Gene names were standardized by replacing underscores with hyphens. Expression values were normalized to Counts Per Million (CPM) using the formula (gene counts / total counts) × 10⁶, followed by a log₂-transformation with a pseudocount of 1. The correlation between modalities was assessed in two contexts: (i) shared genes detected in both modalities; and (ii) all genes expressed in at least one modality, with genes showing zero expression in both modalities excluded from the all-genes analysis. Pearson and Spearman correlation coefficients were calculated for each gene set with corresponding p-values. The gene detection overlap was visualized using a Venn diagram showing modality-specific and shared genes.

### · Clustering quality assessment on downsampled datasets

For each downsampled dataset at both nuclei and sequencing depth level, dimensionality reduction, neighborhood graph construction, and Leiden clustering were performed. Clustering quality was evaluated using K-nearest-neighbor (K-NN) purity, which measures local cluster consistency, and Adjusted Rand Index (ARI), which quantifies agreement between clustering results and the known species labels (human, mouse, dog). UMAP embeddings were generated to visually assess species separation and clustering stability across downsampling replicates. Reported values correspond to the mean across the nine independently generated datasets, while error bars represent standard deviation across sampling and downsampling runs.

### · Proliferation score assessment on downsampled datasets

Proliferation labels were transferred from the full-depth dataset based on cell-cycle phase classification: nuclei classified as S-phase or G2/M-phase were considered proliferating, whereas G1 nuclei were considered non-proliferating. For each dataset, a proliferation score was calculated as the combined S-score and G2/M-score and proliferating and non- proliferating nuclei were compared using the Mann–Whitney U test. Biological signal recovery was quantified using the rank-biserial correlation effect size, together with summary statistics describing the score distributions in each group. Reported values represent the mean across the nine downsampled datasets, and error bars represent the standard deviation across sampling and downsampling runs.

### · Differential expression analysis between NIH-3T3 subclusters

Differential expression analysis was performed at the transcript level, with the focus on comparing the two known mouse subpopulations present within NIH-3T3 cells. First, the pooled isoform-by-cell matrices were subsampled to 3,000 nuclei belonging to either of the two mouse clusters. As in the previous downsampling procedures, sequencing depth was also reduced to 3,000 UMIs/nucleus, yielding nine matrices per platform.

Transcript counts were aggregated by sample and cell cluster using the decoupler library (v1.8.0) (https://doi.org/10.1093/bioadv/vbac016). Only groups with a minimum of 20 nuclei were retained to ensure sufficient statistical power. Lowly expressed genes were filtered using edgeR’s filterByExpr function. Differential expression analysis was conducted using edgeR via the pertpy toolkit. Pairwise comparisons were performed between both cell clusters, with the smallest subpopulation being considered as the baseline. Raw p-values were adjusted for multiple testing using the Benjamini-Hochberg method. Results were filtered to retain only isoforms meeting the following criteria: (i) false discovery rate (FDR) <5%; (ii) |log₂ fold change| >1 (≥2-fold change); and (iii) logCPM >1.0 (average log counts per million to exclude lowly expressed genes).

### · Gene ontology analyses

Gene ontology analyses were performed using ShinyGO v.0.85.1^26^ with the “mmusculus_gene_ensembl” annotation, “Mouse genes GRCm39” assembly (Taxonomy ID 10090, Ensembl) and “GO Biological Process” as pathway database with a FDR cutoff of 0.05.

## Results

To assess the suitability of different snRNA-seq assays for isoform-resolved transcriptomics, cDNA samples were generated in triplicate using the 10x Genomics 3’ and 5’, Parse Biosciences, and ArgenTag assays, respectively, with nuclei extracted from a benchmark sample composed of a mixture of human (HEK293T, 40%), mouse (NIH-3T3, 40%), and dog (MDCK-2, 20%) cell lines (Fig. 1A). The cDNA samples were then prepared for long-read sequencing with the ONT Ligation Sequencing Kit V14 and sequenced on a ONT PromethION 2 (P2) Solo sequencer using two ONT PromethION DNA R10.4.1 flow cells per library (Fig. 1A, Methods), generating between 138 and 412 M reads per sample (Suppl. Tab. 1).

**Figure 1.**
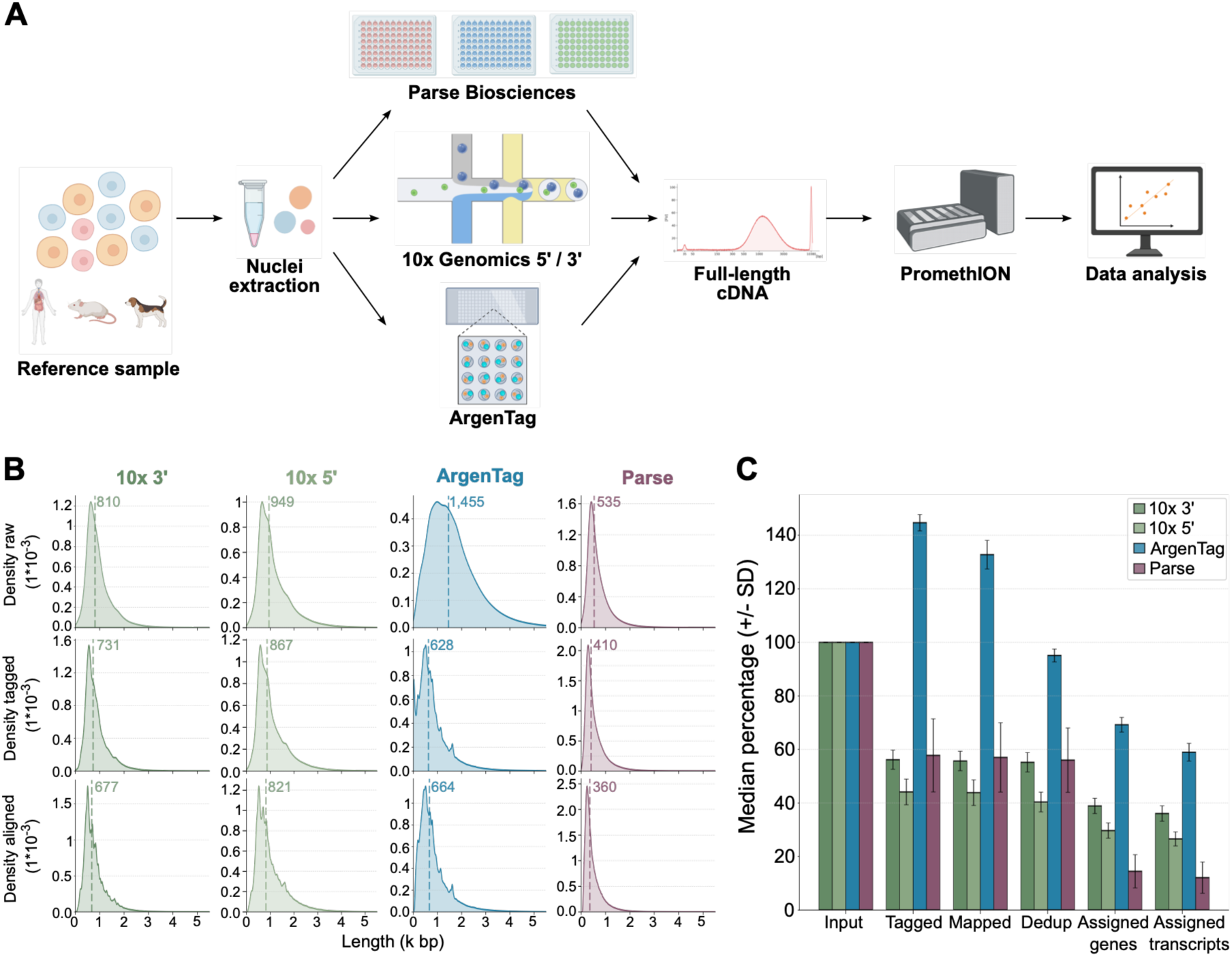
Study overview, raw data quality, and read survival comparison for tested LR snRNA-seq methods. **(A)** Benchmark study overview. Created in BioRender. Kohler, F. (2026) https://BioRender.com/59cxcfu. **(B)** Read length distributions for raw, barcoded (tagged) and aligned reads. **(C)** Read survival defined as the median proportion (%) of input reads (Input) retained after barcode identification (Tagged), mapping (Mapped), deduplication (Dedup), as well as gene (Assigned genes) and isoform (Assigned transcripts) quantification. Error bars represent the standard deviation (SD).

Direct comparison of single-cell RNA sequencing platforms can be confounded by the use of vendor-specific processing pipelines that rely on different alignment, barcode correction and quantification strategies. To enable a fair, methodologically consistent comparison, we thus developed a dedicated bioinformatic pipeline to uniformly process data generated with the three platforms (Methods). Specifically, each dataset was first demultiplexed using the vendor-recommended algorithm to obtain reads tagged with corrected cell barcodes and uncorrected UMIs. Subsequently, all datasets were processed uniformly, allowing gene- and transcript-level quantification to be compared across platforms without confounding platform differences with pipeline-specific biases.

### Read length and read yield differences reflect method-specific library preparation strategies

As shown in Fig. 1B, the longest raw reads were generated for the ArgenTag libraries (Median length: 1,455 bp), with noticeably shorter reads observed for Parse Biosciences (Median length: 535 bp), mirroring a shorter original cDNA population (Suppl. Fig. 1A). 10x Genomics 3’ and 5’ libraries displayed median raw read lengths of 810 bp and 949 bp, respectively, broadly representative of their original cDNA lengths (Suppl. Fig. 1A). After barcode assignment and read mapping, the longest reads aligned to the reference were found in the 10x Genomics 5’ data with a median length of 821 bp, followed by 10x Genomics 3’ and ArgenTag datasets with median read lengths of 677 bp and 664 bp, respectively, while Parse Biosciences data contained the shortest aligned reads (Median length: 360 bp) (Fig. 1B).

To quantify effective read yields, defined as the proportion of initial reads ultimately available for gene and transcript quantification, we assessed read retention during processing, including barcode identification (tagging), mapping, deduplication, as well as gene and isoform assignment, respectively. After tagging, only around 50% of input reads generated from 10x Genomics 3’, 5’ and Parse Biosciences libraries were retained (Fig. 1C), indicating substantial read loss due to incorrect barcode identification. For ArgenTag samples, due to the presence of a large proportion of chimeric reads, the number of barcode-assigned reads increased to 145% of input reads (Fig. 1C). During mapping and deduplication, only a small number of reads were filtered out for 10x Genomics and Parse Biosciences samples, whereas around 8% and 30% of tagged reads were excluded during mapping and deduplication for ArgenTag samples, respectively (Fig. 1C and Suppl. Fig. 1B), suggesting substantial cDNA overamplification. After gene and isoform assignment, the largest proportions of input reads were retained for ArgenTag (68% and 59%), followed by 10x Genomics 3’ (40% and 37%) and 10x Genomics 5’ (29% and 26%) samples. For Parse Biosciences samples, a substantial number of reads could not be assigned to genes or isoforms using IsoQuant, resulting in very low retention of reads after gene (15%) and isoform assignment (12%), respectively (Fig. 1C and Suppl. Fig. 1B).

Together, these findings reinforce the notion that library preparation strategies strongly influence raw data metrics, including read length and effective yield. 10x Genomics and Parse Biosciences assays, originally optimized for short-read sequencing, were characterized by shorter raw reads and substantial read losses during barcode assignment, whereas the native long-read approach utilized by ArgenTag was found to produce longer raw reads and superior read retention for gene and isoform quantification, despite extensive chimera formation and apparent cDNA overamplification.

### Different LR snRNA-seq approaches capture qualitatively distinct full-length transcriptomes

To assess how the methods differ with regard to their ability to capture the full-length transcriptome, we next compared gene body coverage between the methods based on the normalized coverage for the 10% shortest (min. 350 bp) and 10% longest transcripts in the combined reference genome, respectively. Whereas all four methods generated relatively even coverage for the shortest transcripts, the broadest coverage for the longest transcripts was observed in the Parse Biosciences data (Fig. 2A and 2B), likely due to the use of both random and poly(A) priming during reverse transcription. 10x Genomics and ArgenTag data, on the other hand, exhibited noticeable 5’ and 3’ biases, respectively (Fig. 2A and 2B), reflecting their exclusive reliance on end-based priming.

**Figure 2.**
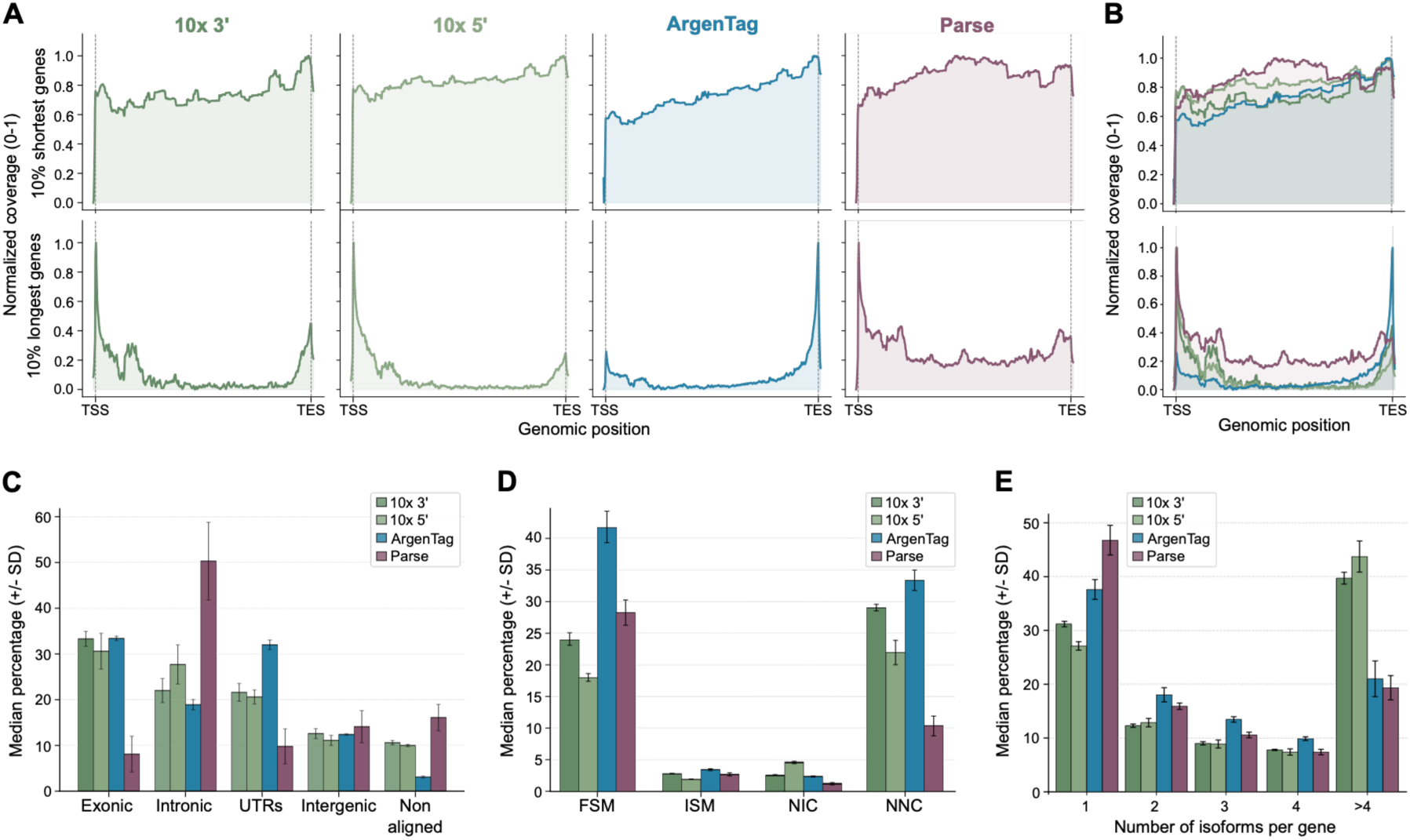
Qualitative characteristics of the long-read transcriptome captured by 10x Genomics 3’ and 5’, ArgenTag, and Parse Biosciences LR snRNA-seq. **(A)** Normalized gene body coverage from Transcription Start Site (TSS) to Transcription End Site (TES) for the 10% shortest (top) and 10% longest (bottom) genes in the combined reference. **(B)** As in (A) but overlaid for all methods. **(C)** Median proportion (%) of bases in exonic, intronic, untranslated (UTRs), intergenic or non-aligned regions. Error bars represent the standard deviation (SD). **(D)** Median proportion (%) of transcripts categorized as full splice match (FSM), incomplete splice match (ISM), novel in catalogue (NIC) or novel not in catalogue (NNC) by SQANTI3^21^. Error bars represent the standard deviation (SD). **(E)** Number of unique transcript isoforms detected per annotated gene. Error bars represent the standard deviation (SD).

Additionally, we analyzed genomic coverage as measured by the percentage of bases in exonic, intronic, untranslated (UTR) or intergenic regions. In the 10x Genomics 3’ and 5’, and ArgenTag data, around 30% of bases mapped to exonic regions, while around 20% of bases were located in intronic regions. Conversely, in the Parse Biosciences data, around 50% of bases covered intronic regions, with only a small proportion of bases derived from exonic regions (9%), again likely related to the use of random priming during reverse transcription (Fig. 2C).

We also assessed the ability of each of the methods to accurately identify known and novel transcript variants using SQANTI3^21^. The highest percentage of known isoforms, i.e. those matching a reference transcript (full splice match), was identified in ArgenTag data (42%), followed by Parse Biosciences (28%) and 10x Genomics 3’ data (23%) (Fig. 2D). In turn, the largest share of new transcript variants with either known (Novel-in-catalogue) or novel splice donors and/or acceptors (Novel-not in catalogue) was found in 10x Genomics (3’: 3% and 29%, 5’: 5% and 23%) and ArgenTag (2% and 33%) datasets, whereas Parse Biosciences data were characterized by substantially smaller shares of novel isoforms and splice junctions (1% and 10%) (Fig. 2D and Suppl. Fig. 2A-C). Interestingly, the methods also differed with regard to the number of transcript isoforms identified per gene, with both 10x Genomics datasets containing a substantially larger proportion of genes with 4 or more isoforms (Fig. 2E), suggesting superior transcript recovery in these assays.

Taken together, these findings demonstrate that the four workflows capture qualitatively distinct transcript populations shaped by their underlying cDNA generation strategies. 10x Genomics and ArgenTag datasets showed enhanced capture of exonic transcript regions and efficient recovery of full-length splice isoforms including novel variants, while Parse Biosciences data preferentially contained shorter, intron-rich and previously characterized transcripts, consistent with its random priming strategy.

### Quantitative differences in isoform detectability despite high concordance with matched short-read snRNA-seq data

As a next step, we compared the four methods regarding their ability to quantify the isoform- resolved transcriptome using IsoQuant^20^. To account for differences in sequencing depth as well as nuclei numbers, the three biological replicates were pooled for every method and 3,000 nuclei were sampled at random, followed by downsampling to different sequencing depths (Methods).

As shown in Fig. 3A, Parse Biosciences data contained the highest aggregate number of genes and isoforms, while the ArgenTag assay yielded lower overall numbers of distinct genes and isoforms, a pattern that persisted even at lower sequencing depths (Suppl. Fig. 3A and B). Conversely, when comparing the methods for their per-nucleus feature recovery potential, i.e. the ability to identify a new transcript or gene for every additional UMI sampled, we observed higher sensitivity for 10x Genomics and ArgenTag, while Parse Biosciences data allowed for the discovery of slightly less new features for every additional UMI sampled (Fig. 3B and Suppl. Fig. 3C). Of note, as a consequence of moderate sequencing depths and data processing constraints, including merging of data from replicates and subsampling, all datasets were characterized by relatively limited saturation levels of around 40–50% at the transcript level and only slightly higher levels (44–51%) at the gene level (Fig. 3B and Suppl. Fig. 3C).

**Figure 3.**
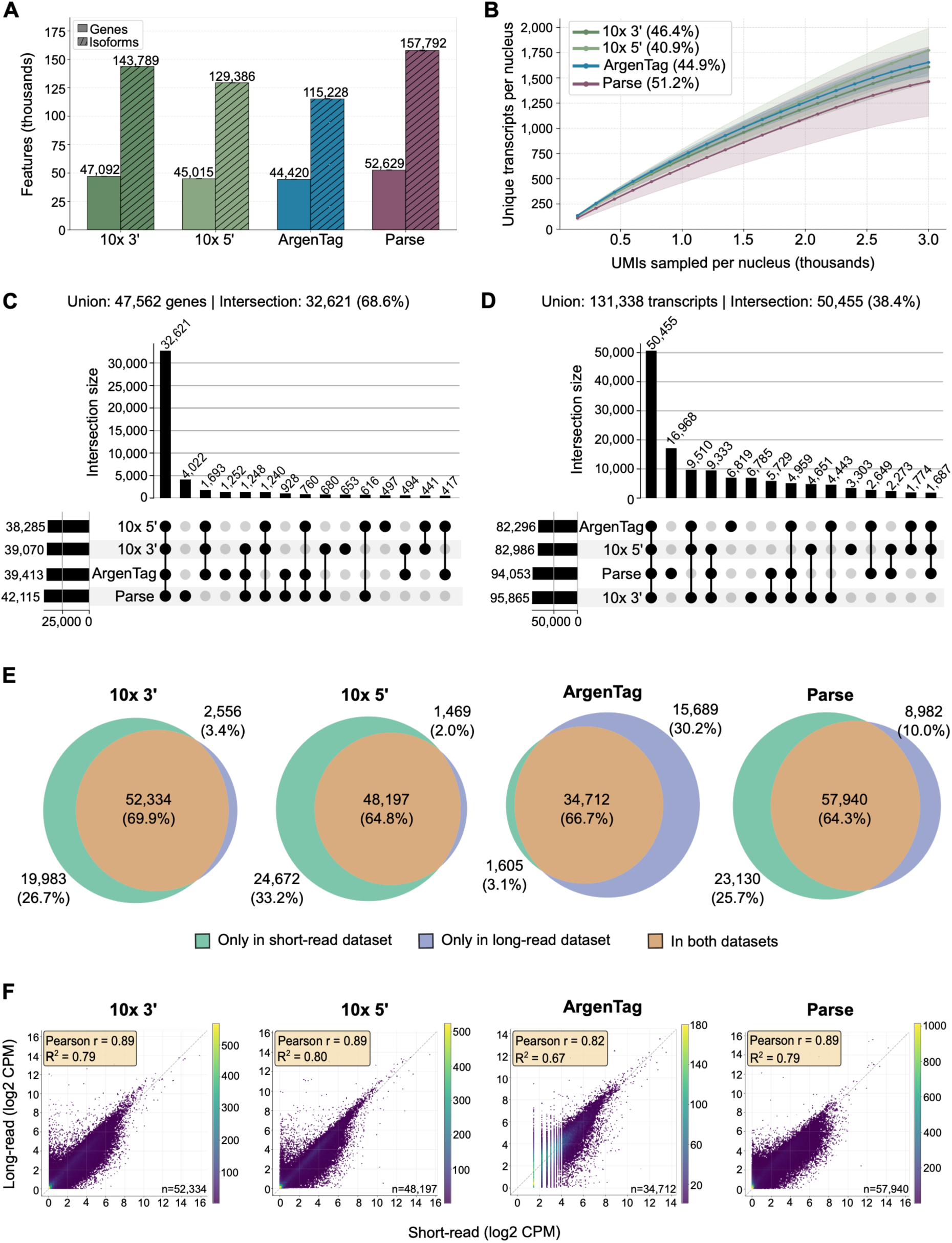
Quantitative characteristics of the long-read transcriptome measured by 10x Genomics 3’ and 5’, ArgenTag, and Parse Biosciences LR snRNA-seq. **(A)** Aggregate numbers of distinct genes and transcript isoforms identified per platform. For each dataset, 3,000 nuclei were selected at random followed by downsampling to 3,000 UMIs/nucleus. **(B)** Per-nucleus sequencing saturation as measured by the number of unique transcripts detected per nucleus for every thousand UMIs sampled. For each dataset, 3,000 nuclei were selected at random. Percentages in brackets indicate saturation levels. **(C)** Overlap of human, mouse, and dog genes detected by one or more platforms in the downsampled matrixes (3,000 nuclei, 3,000 UMIs/nucleus). **(D)** Overlap of human, mouse, and dog transcripts detected by one or more platforms in the downsampled matrixes (3,000 nuclei, 3,000 UMIs/nucleus). **(E)** Overlap of genes detected in short-read (green, left) and long-read (blue, right) data, respectively. Only genes with expression values of >0 counts per million (CPM) were considered. **(F)** Correlation between gene expression values based on long-read (vertical axis) or short-read (horizontal axis) data for those genes detected in both data types. CPM = Counts per million.

Despite their similarities, due to their unique methodological approaches, the four methods could differ with regard to the identity of genes and transcripts detected in a given sample. Consequently, we assessed which genes and transcripts were identified by more than one method. Importantly, the majority of genes (32,621; 68.6% of all genes) identified in the benchmark sample was detected by all assays (Fig. 3C). In contrast, considerable variability existed at the transcript level, with 50,455 transcripts, representing only 38.4% of all isoforms, overlapping between methods (Fig. 3D). In addition, 1,693 (3.6%) genes and 9,510 (7.24%) transcripts were shared across the 10x Genomics 5’, 10x Genomics 3’, and ArgenTag data, respectively, while the Parse Biosciences data contained a substantial fraction of genes (4,022; 8.46%) and transcripts (16,968; 12.92%) exclusively found in this method (Fig. 3C and 3D). These findings are consistent with the conclusion that method-specific differences lead to marked variability in feature detection between the assays, especially at the transcript level, with the additional use of randomers by Parse Biosciences supporting the identification of features distinct from those captured by the other, exclusively poly-A capture-based methods. Short-read sequencing currently represents the gold-standard for the analysis of single-cell and single-nucleus transcriptomes, thanks to an attractive yield-to-cost ratio and lower error rates. Thus, we also analyzed how well long-read sequencing-based approaches recapitulate conventional, short-read sequencing-based single-nucleus transcriptomic readouts. For this purpose, one cDNA sample for every method was prepared for short-read sequencing on the Illumina platform to compare gene-level expression profiles qualitatively and quantitatively (Methods). Of note, in the case of ArgenTag, after the preparation of short-read sequencing libraries, barcodes were detectable in only a fraction of reads, therefore limiting the sequencing depth and discovery power for this assay.

Importantly, the majority of genes, ranging from 64.3% in Parse Biosciences to 69.9% in 10x Genomics 3’, was detected in both long and short-read datasets across all methods (Fig. 3E), indicating that gene-level transcriptomic information is generally well recovered using long-read sequencing. However, a substantial proportion of genes in the 10x Genomics 5’ (33.2%), 10x Genomics 3’ (26.7%) and Parse Biosciences (25.7%) data was exclusively identified in the corresponding short-read datasets, hinting at the potential of additional gene recovery upon deeper long-read sequencing. For ArgenTag, due to the yield constraints observed in the short-read data, the majority of non-overlapping genes were derived from the long-read sequencing data (30.2%) (Fig. 3E), with only 3.1% of genes exclusively found in the short-read data.

Quantitatively, there was generally a strong correlation between expression levels oberserved in the short and long-read sequencing data, respectively, for 10x Genomics and Parse Biosciences, with Pearson correlation coefficients of around 0.89 (Fig. 3F). Conversely, for the ArgenTag assay, owing to the substantially lower sequencing depth in the short-read dataset, a lower concordance of expression values (Pearson r = 0.82) was observed (Fig. 3F).

Together, these results indicate that, after accounting for differences in nuclei numbers and sequencing depth, quantitative performance varies substantially across platforms and depends on the metric examined. Although the Parse Biosciences workflow exhibited slightly lower transcript-level sensitivity and reduced concordance with matched short-read data, it yielded higher overall numbers of detected genes and isoforms, including a substantial set of features not detected by the poly(A)-based methods, underscoring that random priming-based approaches interrogate partially distinct transcript populations. In contrast, 10x Genomics 3’ and 5’ datasets demonstrated superior per-cell feature recovery and stronger correlation with short-read data, supporting their quantitative robustness for gene-level analyses. Nevertheless, at standard sequencing depths, all four methods provide only a relatively sparse representation of the full-length transcriptome, reflecting current throughput-to-cost ratio limitations associated with long-read single-nucleus transcriptomics.

### 10x Genomics 3’, 10x Genomics 5’, Parse Biosciences, and ArgenTag LR snRNA-seq data enable isoform-resolved, single-nucleus transcriptomic analyses

Next, we investigated whether data of all four methods can be used for common single-cell transcriptomic analyses such as unsupervised clustering, differential gene expression, as well as population-specific isoform expression. To do so, we relied on the heterogeneity of the benchmark sample, consisting of 40% human (HEK293T), 40% mouse (NIH-3T3), and 20% dog (MDCK-2) cells. Initially, we assessed whether the methods would correctly quantify the expected proportions of cell types. As shown in Fig. 4A and B, after removing low-quality and mixed nuclei, the smallest deviations from the expected proportions were observed in ArgenTag and Parse Biosciences data, indicating that the initial sample composition was well preserved in these datasets. Conversely, 10x Genomics 3’ and 5’ datasets contained notably smaller numbers of mouse nuclei and substantially higher numbers of human nuclei than expected (Fig. 4A and B). However, we also observed increased proportions of mixed nuclei in some 10x Genomics samples (Suppl. Fig.4A), indicating that problems related to sample processing rather than method-specific capture biases may have accounted for these deviations.

**Figure 4.**
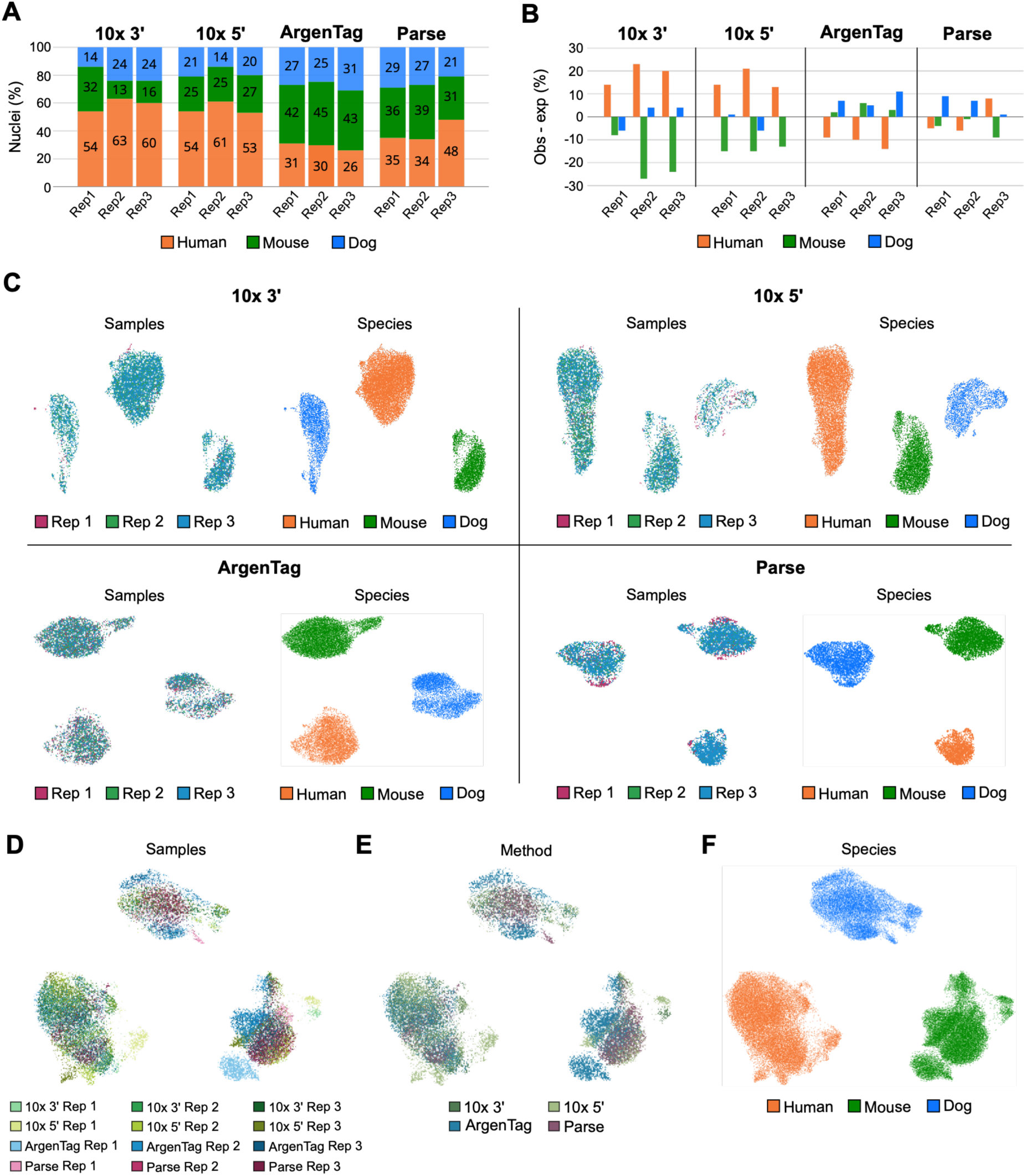
Species assignment, clustering, and integration analyses based on LR snRNA-seq data. **(A)** Sample composition as measured by the proportion (%) of nuclei from human, mouse or dog origin in analyzed replicates (Rep1-3) and **(B)** deviation from the expected distribution (40% human, 40% mouse, and 20% dog). **(C)** Unsupervised clustering of nuclei after removal of low-quality nuclei and batch correction displayed by sample origin (left) and species assignment (right). **(D), (E)** and **(F)** Unsupervised clustering of nuclei after integration of all methods displayed by sample, method, and species, respectively.

Furthermore, we analyzed whether the four methods enable the correct separation of different cell types using unsupervised clustering. Importantly, at maximum sequencing depths, each method successfully resolved the biological heterogeneity of the benchmark sample, identifying distinct human, mouse and dog clusters, as well as a smaller sub-cluster in the NIH-3T3 cell line (Fig. 4C), whose heterogeneity has previously been described^27,28^. We therefore also performed downsampling analyses to evaluate the clustering performance at lower sequencing depths. While all four methods successfully identified species-level differences at higher sequencing depths (>1,000 UMI/nucleus), the ArgenTag assay exhibited slightly inferior performance at lower sequencing depths (<1,000 UMI/nucleus) (Suppl. Fig. 4B–D).

To evaluate potential methodological biases, we next assessed whether LR snRNA-seq data from all four platforms can be integrated, enabling the separation of nuclei based on their species of origin rather than technical sources. As shown in Fig. 4D-F, after integration and batch correction, nuclei clustered by their species of origin rather than by sample identity or method, indicating that integration of data generated by the different LR snRNA-seq methods is possible while preserving species-level biological complexity.

Finally, we aimed to demonstrate the potential of isoform-resolved snRNA-seq data for differential expression analyses at the single-nucleus level. The NIH-3T3 cell line serves as a useful model for this purpose, as it is derived from a whole-mouse embryo and its cellular heterogeneity, including diverse cell types of the fibroblast lineage, is well established^27,28^. Thus, using edgeR^29^, we performed differential isoform expression analyses between the two NIH-3T3 subclusters after downsampling (Methods). Interestingly, while the number of differentially expressed isoforms identified by the individual assays ranged from 1,567 in the 10x Genomics 5’ data to 2,277 in the ArgenTag data (Suppl. Fig. 5A), 99 transcripts were consistently detected by all four methods (Fig. 5A), most of which showed increased expression in the smaller NIH-3T3 subcluster (Fig. 5B). These transcripts were strongly enriched for mitotic and cell cycle markers (Suppl. Fig. 5B), indicating that distinct proliferative programs detectable by all methods underlie NIH-3T3 heterogeneity. Additional sets of differentially expressed isoforms were shared between the ArgenTag and 10x Genomics datasets, but not Parse Biosciences (Fig. 5A and B), suggesting that increased per-cell sensitivity in these assays facilitates the reconstruction of additional isoform-specific expression patterns underlying NIH-3T3 heterogeneity. Importantly, across all methods, cell cycle scoring at the transcript level confirmed that the existence of a proliferating and a non- proliferating cluster underpins the observed NIH-3T3 heterogeneity, with both G2/M and S-phase scores enriched in the smaller subcluster (Fig. 5C). However, after accounting for nuclei numbers and sequencing depth differences, we noticed that this biological signal was best preserved in the ArgenTag data (Suppl. Fig. 5C).

**Figure 5.**
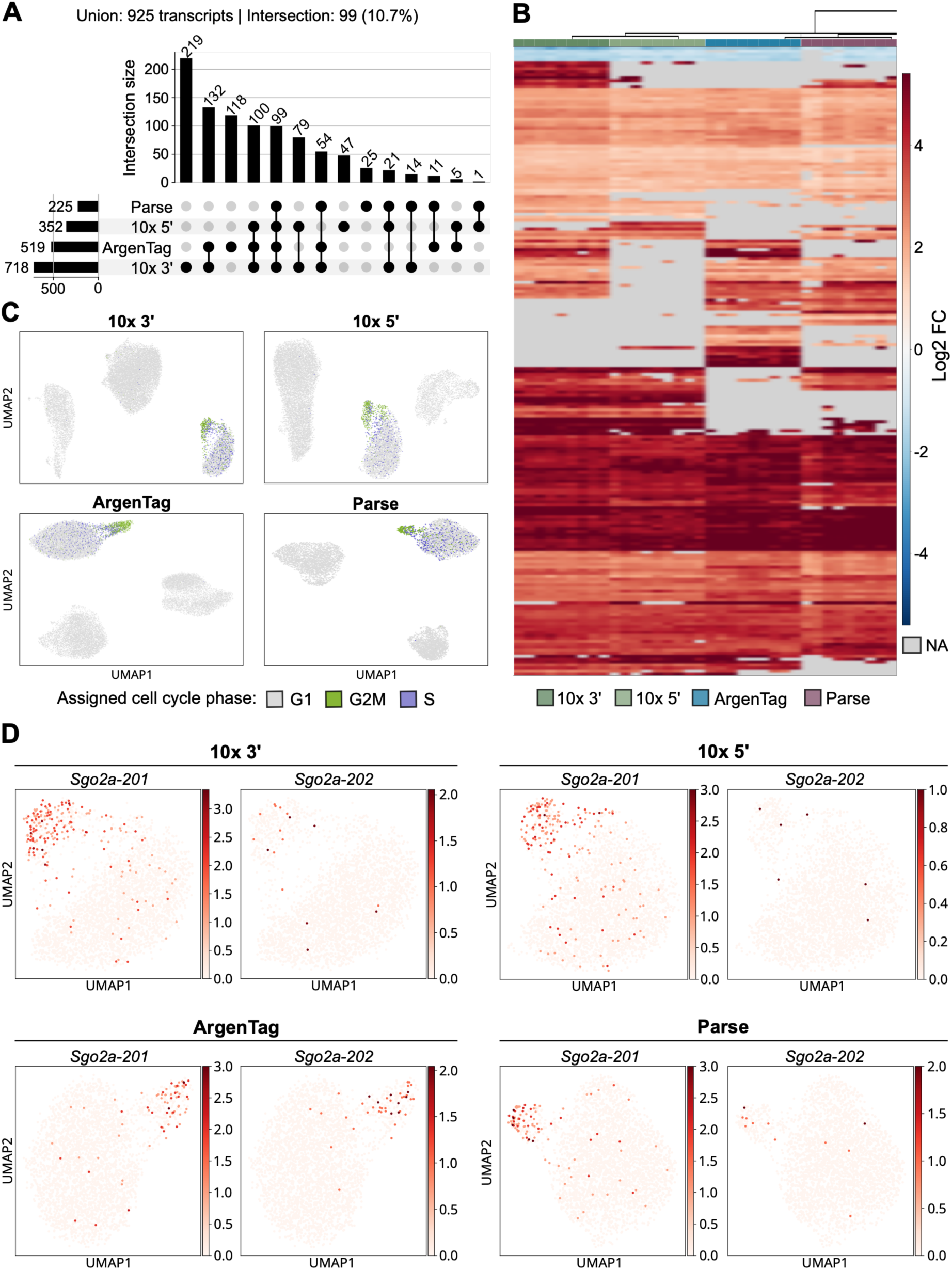
Differential isoform expression analysis in NIH-3T3 nuclei. **(A)** Overlap of transcripts detected as differentially expressed between NIH-3T3 subclusters by one or more methods. **(B)** Expression changes (Log2FC) for isoforms identified as differentially expressed between NIH-3T3 subclusters. Columns represent experiments (n=36), with 9 experiments per method.

At the single-nucleus level, as a proof of concept, we investigated whether the expression of a gene of interest could be attributed to a specific isoform, and whether such isoform-specific expression could be detected by all methods. *Shugoshin 2A* (*Sgo2a*), one of the genes for which one or more transcripts were found to be upregulated in the smaller NIH-3T3 subcluster across all methods, encodes a regulatory protein contributing to accurate chromosome segregation during cell division^30^. According to the Ensembl database^31^, two isoforms are derived from *Sgo2a*: a protein-coding variant, *Sgo2a-201* (ENSMUST00000027202), and a retained-intron variant, *Sgo2a-202* (ENSMUST00000190746). We thus analyzed expression patterns of both transcripts in the NIH-3T3 subcluster, as well as their differential detection by the four assays. Importantly, increased expression of *Sgo2a-201* was observed in the smaller NIH-3T3 subcluster in all datasets (Fig. 5D and Suppl. Tab. 6). In contrast, *Sgo2a-202* was identified as upregulated in the ArgenTag and 10x Genomics 3’ datasets, but not 10x Genomics 5’ and Parse Biosciences datasets (Fig. 5D and Suppl. Tab. 6), indicating substantial method- specific variability regarding isoform detectability.

Collectively, these findings demonstrate that all four methods preserve sufficient biological information for standard single-cell transcriptomic analyses, including unsupervised clustering, dataset integration, and isoform-specific differential expression analyses. While all assays captured the biological heterogeneity of our benchmark sample, including the existence of two NIH-3T3 subclusters with distinct proliferative patterns, the isoform-level analysis revealed specific, method-dependent differences in transcript sensitivity, with the ArgenTag and 10x Genomics assays enabling a more granular characterization of subcluster- specific isoform expression.

Rows represent isoforms. NA = not differentially expressed. **(C)** Assigned cell cycle phases for NIH-3T3 subpopulations. As murine cell cycle marker genes were used, human and dog nuclei are expected to be assigned to the G1 phase. **(D)** Expression of *Sgo2a-201* (ENSMUST00000027202, left) and *Sgo2a-202* (ENSMUST00000190746, right) isoforms in 10x Genomics 3’, 10x Genomics 5’, ArgenTag, and Parse Biosciences data.

## Discussion

Long-read single-cell and single-nucleus transcriptomic studies hold tremendous potential for studying transcriptomic diversity in health and disease. However, their wider adoption has been hindered by technological challenges such as limited throughput, elevated error rates compared to short-read alternatives, a lack of suitable processing pipelines, and high assay costs. Consequently, there is currently no broadly established standard approach for LR sc/snRNA-seq, highlighting the need for a systematic benchmarking study to identify methodological trade-offs and guide protocol selection for different biological applications. Here, we present a comprehensive assessment of four current options for performing LR snRNA-seq. Using a standardized benchmark sample composed of human, mouse, and dog nuclei paired with long-read sequencing on the ONT platform, we analyzed raw data quality, transcriptomic coverage and sensitivity, correlation with short-read data, and the suitability for isoform-resolved analyses at the single-nucleus level.

Overall, our results demonstrate that the 10x Genomics 3’ and 5’ assays currently provide the most balanced performance for LR snRNA-seq studies on the ONT platform. Despite substantial read losses during barcode assignment – a common challenge when adapting sc/snRNA-seq workflows originally developed for short-read sequencing to long-read platforms with reduced raw read accuracy^14^ – both assays exhibited strong qualitative and quantitative performance. This was reflected by longer transcript lengths, broad exon coverage, and the effective capture of known and novel splice isoforms, alongside the best quantitative performance among the tested methods in terms of per-cell sensitivity and concordance with matched short-read sequencing data. Both assays also effectively resolved the species heterogeneity in our benchmark sample and preserved NIH-3T3 cluster-specific biological signals. Collectively, these findings are in line with previous studies demonstrating the suitability of the 10x Genomics assays for isoform-resolved single-cell and single-nucleus transcriptomics^7–13^. Whether meaningful differences exist between the 10x Genomics 3’ and 5’ assay versions regarding their suitability for isoform-resolved single-cell and single-nucleus transcriptomics remains an open question. A recent study reported a superior read length profile and more exonic UMIs in the 5’ version relative to the 3’ alternative^32^. Interestingly, these observations were only partially recapitulated in our dataset. Although the 5’ assay yielded slightly longer transcripts and a higher per-cell feature recovery, both assay versions produced broadly comparable results across most evaluated metrics and provided the best overall performance of the methods assessed in this work, thus underscoring their general suitability for LR sc/snRNA-seq studies.

To our knowledge, this study provides the first comprehensive assessment of the ArgenTag assay’s performance in the context of isoform-resolved single-nucleus transcriptomics. It produced the longest raw reads and the highest effective read retention among all tested methods, although the presence of read chimeras artificially inflated raw read lengths and resulted in shorter effective read lengths than in the 10x Genomics datasets. Its qualitative performance was largely comparable to that of the 10x Genomics 3’ and 5’ assays, with a broad feature coverage and effective detection of known and novel isoforms. However, it underperformed quantitatively, yielding lower aggregate numbers of genes and transcripts and the lowest correlation with matched short-read sequencing data. Species separation was possible at higher sequencing depths, while some of the NIH-3T3-specific biological signals were best preserved across all tested methods. Our findings thus underscore the assay’s suitability for isoform-resolved sc/snRNA-seq studies and demonstrate its considerable potential, although further optimization may be required to reduce chimera formation and cDNA overamplification.

In contrast, the Parse Biosciences assay exhibited a performance profile that may make it more suitable for LR sc/snRNA-seq applications where broad transcript recovery and cost efficiency are required. Its random priming strategy likely led to the preferential capture of shorter, intron-enriched transcripts, resulting in a qualitatively distinct transcript population. Nevertheless, the assay demonstrated robust performance, yielding the highest aggregate number of genes and transcripts identified in our samples and good concordance with matched short-read sequencing data, despite a lower per-cell feature discovery power. These findings are consistent with a recent study highlighting the suitability of the Evercode assay for LR scRNA-seq, owing to its ability to capture complementary transcript populations, including non-coding variants, as well as its attractive cost efficiency and multiplexing capacity^33^. Together, these findings support the use of Parse Biosciences as an alternative strategy for LR sc/snRNA-seq studies, particularly when scalability and the recovery of additional RNA populations are priorities.

Importantly, all tested methods preserved sufficient biological information for robust species separation, dataset integration, and isoform-level analyses at single-nucleus resolution, underscoring the overall maturity of current LR snRNA-seq approaches and highlighting the distinct trade-offs associated with different experimental strategies.

It is worth noting that, beyond the instructions provided in the ONT protocol for the Ligation Sequencing Kit V14 and minor adaptations to the ArgenTag workflow, no optimizations of the library preparation workflow – such as fragment size selection, targeted amplification, or exome enrichment – were performed. Designed to enrich cDNA libraries for intact and longer fragments, or to target coding transcripts in particular, such approaches have previously been shown to improve sequencing efficiency and transcriptome quality^7,8,34^. Although beyond the scope of this work, these optimizations represent promising avenues for future LR snRNA-seq studies.

While this benchmark constitutes a comprehensive comparison of the practical performance of current LR snRNA-seq approaches, thereby providing a reliable reference for future isoform-resolved single-nucleus studies, several limitations should be considered when interpreting the findings presented herein. First, the use of single nuclei instead of whole cells represents both a practical advantage and a biological constraint. Single-nuclei approaches are better suited for frozen tissues and difficult-to-dissociate samples, and are therefore especially relevant for many clinical and translational applications, as well as organism-level atlasing efforts. In the context of technical benchmarking of long-read sequencing-based approaches, the use of nuclei offers unique advantages and challenges for transcriptome analysis. On the one hand, long reads can capture incompletely processed transcripts, retained introns, and nascent RNA species within their native nuclear context, enabling the analysis of RNA maturation and splicing, which is difficult to achieve using short-read sequencing-based approaches. On the other hand, nuclear RNA differs substantially from whole-cell RNA, being enriched for incompletely processed and intron-containing transcripts, and the performance differences observed in this study may therefore not fully translate to single-cell protocols using cytoplasmic RNA. In particular, the elevated intronic signal observed in the Parse Biosciences data may represent a direct consequence of using random priming within the nuclear environment.

Second, a key constraint of this study is a relatively low sequencing depth, resulting in modest per-cell saturation levels of around 50%. While this reflects current throughput-to-cost ratio constraints of long-read sequencing platforms, our analyses primarily characterized method performance within the observed depth range rather than providing reliable estimates of maximum achievable transcript complexity. Reaching saturation levels comparable to short-read sequencing-based benchmarks would require substantially greater sequencing investments – resources that were beyond the scope of this study – and represents a challenge across the long-read single-cell sequencing field that will hopefully be addressed by future technological advancements. Similarly, limited nuclei numbers within individual replicates required pooling prior to analysis. This enabled meaningful downsampling analyses but precluded the formal estimation of replicate-to-replicate variability and a statistical comparison of platform performance. Nevertheless, these analyses address the practical question of information recovery under realistic experimental conditions rather than theoretical replicate-level statistics.

Third, although our benchmark sample, consisting of cultured human, mouse, and dog cell lines, provided a highly controlled experimental framework that enabled rigorous cross-method comparisons and species-mixing validation, cultured cell lines represent a relatively homogenous and biologically simplified system compared to primary tissues. Hence, this study does not address challenges associated with complex tissue architecture, low-input samples, rare cell populations or highly heterogeneous transcriptomes. Future benchmarking studies using primary tissues or clinically relevant samples will therefore be essential to determine how these workflows perform in more complex biological scenarios.

Altogether, our study demonstrates that current commercial LR snRNA-seq workflows enable robust transcriptome analyses at single-nucleus resolution but also exhibit important methodological trade-offs. The 10x Genomics 3’ and 5’ assays appear particularly well suited for comprehensive isoform-resolved transcriptomics, whereas the Parse Biosciences solution may provide complementary information on non-coding and immature RNA populations. The ArgenTag assay emerged as a promising alternative for LR snRNA-seq, although further optimization may be required to maximize its performance. Ultimately, these findings provide a valuable framework for selecting an appropriate workflow for future isoform-resolved transcriptomic studies with cellular resolution.

## Author information

F.K. and B.F.P. jointly conceived the study. Experiments were performed by M.S.A., L.Z.J., M.Z., and F.K. Bioinformatic processing and analyses were performed by A.D.T., L.C. and J.P.. The manuscript was prepared by F.K. and A.D.T. with additional input by all authors. Corresponding author: F.K..

## Acknowledgements

We acknowledge the support of the Spanish Ministry of Science and Innovation through the Centro de Excelencia Severo Ochoa (CEX2020-001049-S, MCIN/AEI/10.13039/501100011033), and the Generalitat de Catalunya through the CERCA programme. Additionally, we thank Joaquin Expeleta (ArgenTag Corp.) for developing and providing custom code that enabled key aspects of the data processing and analysis performed in this study. Finally, we are grateful to the CRG Core Technologies Programme for their support and assistance with this study.

## References

1. Wegener, M. & Müller-McNicoll, M. Nuclear retention of mRNAs – quality control, gene regulation and human disease. Semin. Cell Dev. Biol. 79, 131–142 (2018).

2. Schmid, M. & Jensen, T. H. Controlling nuclear RNA levels. Nat. Rev. Genet. 19, 518– 529 (2018).

3. Joglekar, A., Foord, C., Jarroux, J., Pollard, S. & Tilgner, H. U. From words to complete phrases: insight into single-cell isoforms using short and long reads. Transcription 14, 92–104 (2023).

4. Kumari, P. et al. Advances in long-read single-cell transcriptomics. Hum. Genet. 143, 1005–1020 (2024).

5. Wang, F., Xing, Y. & Lin, L. Beyond gene expression: Single-cell transcriptomics at isoform resolution. Trends in Genetics (2026) doi:10.1016/j.tig.2026.05.013.

6. De Simone, M. et al. A comprehensive analysis framework for evaluating commercial single-cell RNA sequencing technologies. Nucleic Acids Res. 53, (2025).

7. Hardwick, S. A. et al. Single-nuclei isoform RNA sequencing unlocks barcoded exon connectivity in frozen brain tissue. Nat. Biotechnol. 40, 1082–1092 (2022).

8. Joglekar, A. et al. Single-cell long-read sequencing-based mapping reveals specialized splicing patterns in developing and adult mouse and human brain. Nat. Neurosci. 27, 1051–1063 (2024).

9. Philpott, M. et al. Nanopore sequencing of single-cell transcriptomes with scCOLOR-seq. Nat. Biotechnol. 39, 1517–1520 (2021).

10. Tian, L. et al. Comprehensive characterization of single-cell full-length isoforms in human and mouse with long-read sequencing. Genome Biol. 22, 310 (2021).

11. Rebboah, E. et al. Mapping and modeling the genomic basis of differential RNA isoform expression at single-cell resolution with LR-Split-seq. Genome Biol. 22, 286 (2021).

12. Lebrigand, K., Magnone, V., Barbry, P. & Waldmann, R. High throughput error corrected Nanopore single cell transcriptome sequencing. Nat. Commun. 11, 4025 (2020).

13. Gupta, I. et al. Single-cell isoform RNA sequencing characterizes isoforms in thousands of cerebellar cells. Nat. Biotechnol. 36, 1197–1202 (2018).

14. Ezpeleta, J. et al. Robust and scalable barcoding for massively parallel long-read sequencing. Sci. Rep. 12, 7619 (2022).

15. Trull, A., Worthey, E. A. & Ianov, L. scnanoseq: an nf-core pipeline for Oxford Nanopore single-cell RNA-sequencing. Bioinformatics 41, (2025).

16. You, Y. et al. Identification of cell barcodes from long-read single-cell RNA-seq with BLAZE. Genome Biol. 24, 66 (2023).

17. Li, H. New strategies to improve minimap2 alignment accuracy. Bioinformatics 37, 4572–4574 (2021).

18. Kinsella, R. J. et al. Ensembl BioMarts: a hub for data retrieval across taxonomic space. Database 2011, bar030–bar030 (2011).

19. Smith, T., Heger, A. & Sudbery, I. UMI-tools: modeling sequencing errors in Unique Molecular Identifiers to improve quantification accuracy. Genome Res. 27, 491–499 (2017).

20. Prjibelski, A. D. et al. Accurate isoform discovery with IsoQuant using long reads. Nat. Biotechnol. 41, 915–918 (2023).

21. Pardo-Palacios, F. J. et al. SQANTI3: curation of long-read transcriptomes for accurate identification of known and novel isoforms. Nat. Methods 21, 793–797 (2024).

22. Ramírez, F. et al. deepTools2: a next generation web server for deep-sequencing data analysis. Nucleic Acids Res. 44, W160–W165 (2016).

23. Wolf, F. A., Angerer, P. & Theis, F. J. SCANPY: large-scale single-cell gene expression data analysis. Genome Biol. 19, 15 (2018).

24. Wolock, S. L., Lopez, R. & Klein, A. M. Scrublet: Computational Identification of Cell Doublets in Single-Cell Transcriptomic Data. Cell Syst. 8, 281–291.e9 (2019).

25. Tirosh, I. et al. Dissecting the multicellular ecosystem of metastatic melanoma by single-cell RNA-seq. Science (1979). 352, 189–196 (2016).

26. Ge, S. X., Jung, D. & Yao, R. ShinyGO: a graphical gene-set enrichment tool for animals and plants. Bioinformatics 36, 2628–2629 (2020).

27. Xu, K. & Rubin, H. Cell transformation as aberrant differentiation: Environmentslly dependent spontaneous transformation of NIH 3T3 cells. Cell Res. 1, 197–206 (1990).

28. Rahimi, A. M., Cai, M. & Hoyer-Fender, S. Heterogeneity of the NIH3T3 Fibroblast Cell Line. Cells 11, 2677 (2022).

29. Robinson, M. D., McCarthy, D. J. & Smyth, G. K. edgeR: a Bioconductor package for differential expression analysis of digital gene expression data. Bioinformatics 26, 139–140 (2010).

30. Yao, Y. & Dai, W. Shugoshins function as a guardian for chromosomal stability in nuclear division. Cell Cycle 11, 2631–2642 (2012).

31. Dyer, S. C. et al. Ensembl 2025. Nucleic Acids Res. 53, D948–D957 (2025).

32. Hsu, J. et al. Comparing 10x Genomics single-cell 3’ and 5’ assay in short-and long-read sequencing. Preprint at 10.1101/2022.10.27.514084 (2022).

33. Xu, Z. et al. SCOTCH: isoform-level characterization of gene expression through long-read single-cell RNA sequencing. Nat. Commun. (2026) doi:10.1038/s41467-026-72665-5.

34. Michielsen, L. et al. Spatial isoform sequencing at sub-micrometer single-cell resolution reveals novel patterns of spatial isoform variability in brain cell types. Preprint at 10.1101/2025.06.25.661563 (2025).

