## Supplementary Material for "Systematic benchmarking of commercial workflows for isoform-resolved single-nucleus transcriptomics"

**Author information:**

[1]Centre for Genomic Regulation (CRG), The Barcelona Institute of Science and Technology, Dr. Aiguader 88, Barcelona, 08003, Spain.
[2]Universitat Pompeu Fabra (UPF), Barcelona, Spain.

**Corresponding author:**

Florian Köhler
Centre for Genomic Regulation (CRG)
C/ Dr. Aiguader 88, Barcelona, 08003, Spain

https://orcid.org/0000-0002-8650-9574

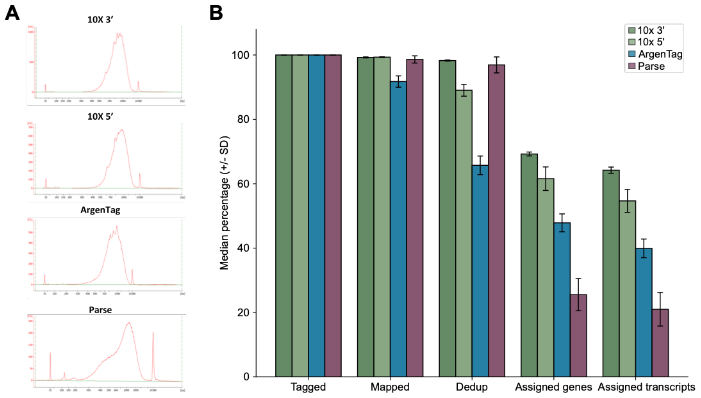


**Suppl. Figure 1. Comparison of cDNA profiles and post-tagging read survival across evaluated methods. (A)** Representative Bioanalyzer profiles of cDNA samples derived from the 10x Genomics 3', 10x Genomics 5', ArgenTag, and Parse Biosciences assays. **(B)** Read survival defined as the median proportion (%) of tagged reads (Tagged) retained after mapping (Mapped), deduplication (Dedup), as well as gene (Assigned genes) and isoform (Assigned transcripts) quantification. Error bars represent the standard deviation (SD).

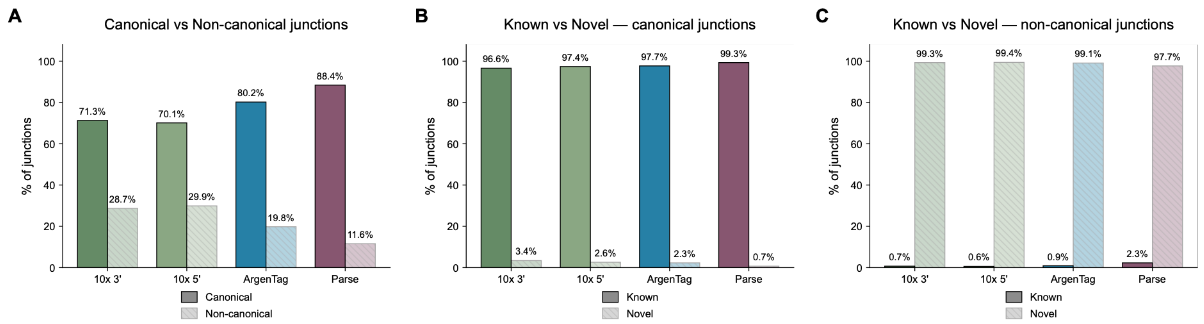


**Suppl. Figure 2. Splice junction detection in 10x Genomics 3’, 10x Genomics 5’, ArgenTag, and Parse Biosciences data.** Proportions of **(A)** canonical vs. non-canonical, **(B)** known vs. novel canonical, and **(C)** known vs. novel non-canonical splice junctions detected in all replicates using SQANTI3[21].

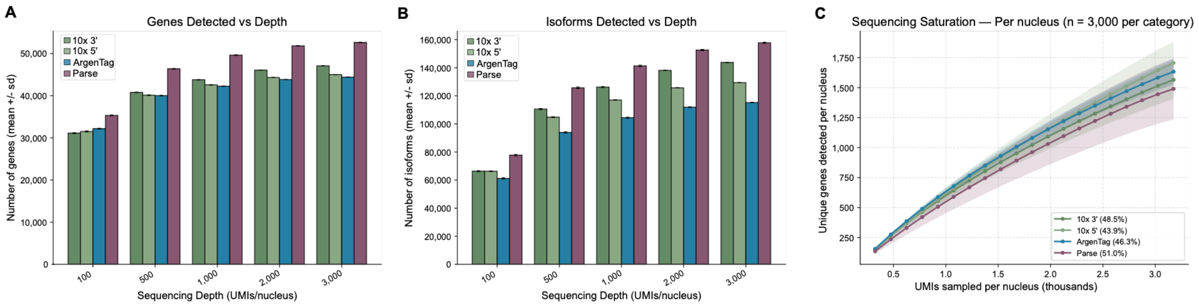


**Suppl. Figure 3. Distinct genes and isoforms detected at different sequencing depths across evaluated methods. (A)** Total number of distinct genes and **(B)** isoforms detected for all three species using IsoQuant[20] after randomly selecting 3,000 nuclei from the merged dataset (replicates). For each platform and cell-number condition, nuclei were randomly sampled three independent times, and each sampled dataset was downsampled to the target UMI depth three times. Consequently, the shown values correspond to the mean of nine independent platform-depth measurements (3 cell-sampling replicates × 3 count-downsampling replicates). Error bars represent the associated standard deviation. **(C)** Per-nucleus sequencing saturation as measured by the number of unique genes detected per nucleus for every thousand UMIs sampled. For each dataset, 3,000 nuclei were selected at random. Percentages in brackets indicate saturation levels.

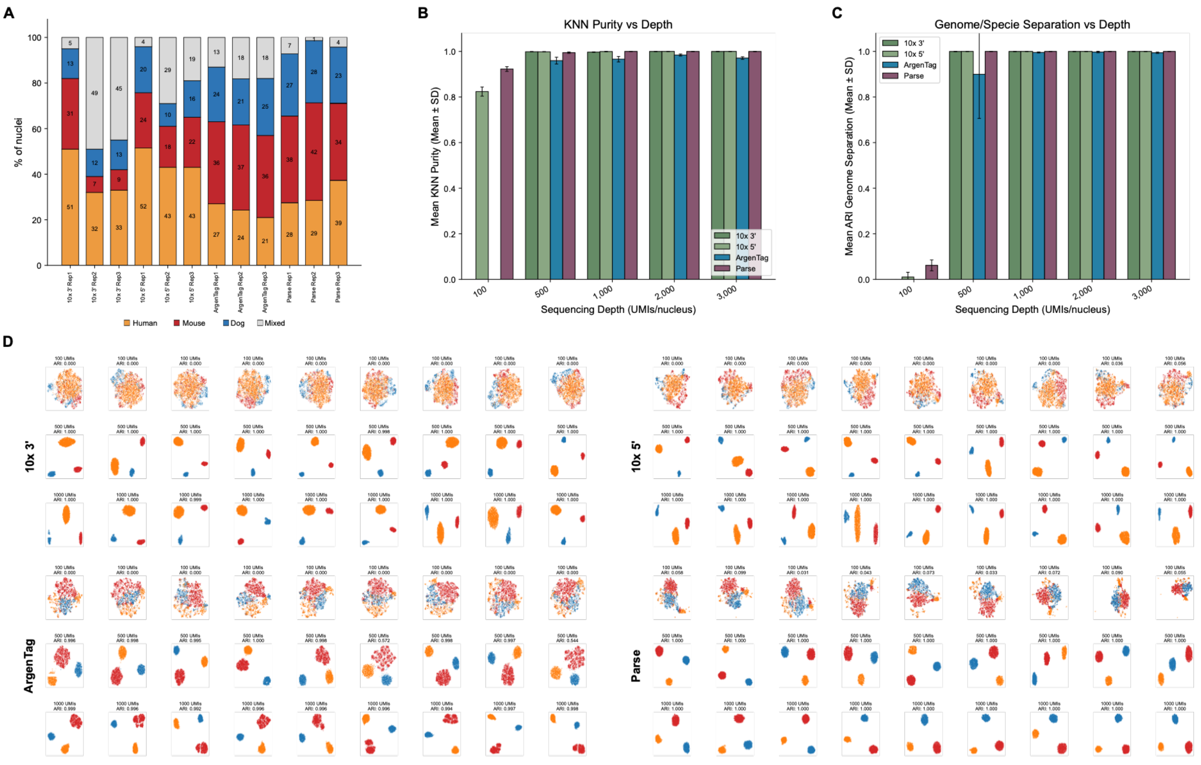


**Suppl. Figure 4. Sample composition, clustering quality, and cluster identity analysis. (A)** Sample composition as measured by the proportion (%) of nuclei from human, mouse, dog, or mixed origin in analyzed replicates. Nuclei were defined as mixed if >20% of the counts align to two or more genomes. **(B)** Mean KNN purity at different sequencing depths. KNN purity is a measure of local neighborhood consistency of cell type assignments (Methods), with values of <0.6 suggesting poorly defined clusters or high inter-cluster mixing, and higher values of 0.8–1.0 indicating well-defined, locally homogeneous clusters. **(C)** Mean Adjusted Rand Index (ARI) at different sequencing depths. The ARI quantifies the agreement between clustering results and the known species labels (Methods). **(D)** Unsupervised clustering at different sequencing depths. Genome-level UMAP plots were generated for 3,000 randomly selected nuclei at the sequencing depths of 100 UMIs/cell (top), 500 UMIs/cell (center), and 1,000 UMIs/cell (bottom).

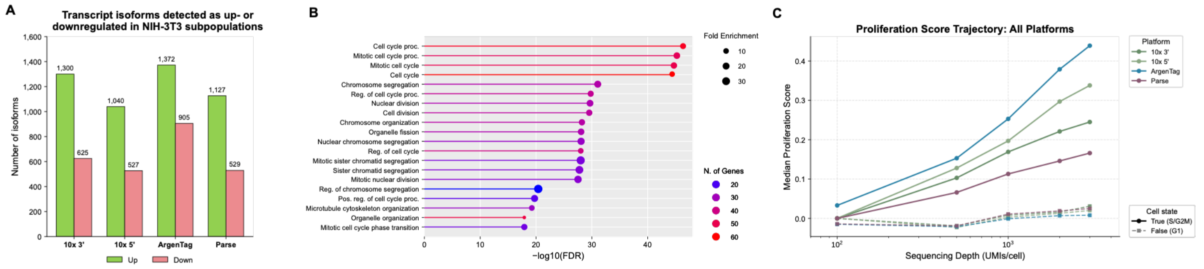


**Suppl. Figure 5. Differential isoform expression analysis in NIH-3T3 nuclei. (A)** Number of transcript isoforms identified as differentially expressed (upregulated/downregulated) in the smaller NIH-3T3 cluster using edgeR[29]. **(B)** Gene ontology (GO) analysis ("GO Biological Process") of 99 transcripts identified as differentially expressed between the two NIH-3T3 subclusters by all methods. "N. of Genes" = Number of Genes. **(B)** Proliferation score at different sequencing depths (UMIs/cell). The score was calculated as the rank-biserial correlation effect size for every dataset, based on the cell cycle phase classification as proliferating (S+G2/M phase) or non-proliferating (G1 phase) (Methods).
